# Structural basis for the neurotropic AAV9 and the engineered AAVPHP.eB recognition with cellular receptors

**DOI:** 10.1101/2022.01.23.477411

**Authors:** Guangxue Xu, Ran Zhang, Huapeng Li, Kaixin Yin, Xinyi Ma, Zhiyong Lou

**Affiliations:** MOE Key Laboratory of Protein Science & Collaborative Innovation Center of Biotherapy, School of Medicine, Tsinghua University, Beijing, China; School of Life Sciences, Tsinghua University, Beijing, China; PackGene Biotech, Guangzhou, Guangdong, China; International School of Beijing, Beijing, China; Beijing No.8 High School, Beijing, China

**Keywords:** AAV9, AAVPHP.eB, AAVR, cryo-EM

## Abstract

Clade F adeno-associated virus (AAV) 9 has been utilized as therapeutic gene delivery vector, and it is capable of crossing blood brain barrier (BBB). Recently, an AAV9 based engineering serotype with enhanced BBB crossing ability, AAVPHP.eB, further expand clade F AAVs’ usages in the central nervous system (CNS) gene delivery. In this study, we determined the cryo-electron microscopy (cryo-EM) structures of the AAVPHP.eB, and its parental serotype AAV9 alone or in complex with their essential receptor Adeno-associated virus receptor (AAVR). These structures reveal the molecular details of their AAVR recognition, where the polycystic kidney disease (PKD) repeat domain 2 (PKD2) of AAVR interact to the 3-fold protrusions and the raised capsid regions between the 2- and 5-fold axes termed the 2/5-fold wall of both AAV9 and AAVPHP.eB virions. The interacting patterns of AAVR to AAV9 and AAVPHP.eB are similar with what was observed in AAV1/AAV2-AAVR complexes. Moreover, we found that AAVPHP.eB variable region VIII (VR-VIII) may independently facilitate the new receptor recognition responsible for enhanced CNS transduction. Our study provides insights into different receptor recognition for engineered AAVPHP.eB and parental serotype AAV9, and further reveal the potential molecular basis underlying their different tropism.

## Introduction

Gene therapy offers a promising therapeutic approach for genetic disorders. With the rapid development of gene delivery vectors, deliver methods play a crucial role in gene therapy. Among various gene therapies deliver vectors, Adeno-associated viruses (AAVs) fulfill the criteria for being highly efficient and non-pathogenic as a viral vector in human body. To date, there have been several FDA approved AAV based gene therapy, and numerous ongoing clinical trials^1–3^.

Naturally occurring serotypes of AAVs have been demonstrated to have varied tropism and transduction efficiencies in tissues^4^, however, therapeutic delivery through the blood brain barrier (BBB) remains a challenge for the application of gene therapy in neurological disorder^5^. Several in vivo selected AAV capsids derived from AAV9, such as AAVPHP.eB and AAVPHP.V1 shows enhanced BBB penetrating ability via intravenous administration^6–8^. These variants have paved the way for precise and non-invasive gene therapy delivery. Adeno-associated virus receptor (AAVR) is a transmembrane glycosylated protein containing five polycystic kidney disease (PKD) extracellular domains^9^. AAVR is reported to likely play a role in AAV tropism, and where AAVs adopt distinct interaction pattern to different PKD domains of AAVR^10,11^. With the assistance of bioinformatic approaches, recent studies have identified a glycosylphosphatidylinositol (GPI) -anchored protein expressed on brain endothelial cells called lymphocyte antigen 6 complex, locus A (LY6A, also known as stem cell antigen-1 [SCA-1]) LY6A to be the cellular receptor responsible for the enhanced movement across BBB^12, 13^.

In this work, we report the cryo-EM structure of engineered AAVPHP.eB and its parental serotype AAV9 as well as their complexes with the universal AAV receptor AAVR. We also explore the interaction between AAVPHP.eB and recently identified receptor LY6A. The structures of AAV with its receptors inform that the 7-amino-acid (TLAVPFK) insertion in AAV capsid variable region VIII (VR-VIII) facilitates AAVPHP.eB with different receptor binding ability independent of conventional AAVR. The receptor interacting residues of engineered AAVPHP.eB also reveal underlying molecular mechanism of its enhanced BBB penetration and CNS transduction.

## Results

### Overall structure of AAV9, AAV-PHP.eB in their native form and AAVR bounded form

We first solved the structure of native AAV9 and AAV-PHP.eB by cryoelectron microscopy at the resolution of 3.87 Å and 2.85 Å at a 0.143 cutoff of FSC **(Supplementary Table 1, Supplementary Figure 1)**. AAV9 and AAV-PHP.eB share common structural features with reported AAV9 crystal structure and other structures of AAV serotypes, including protrusions surrounding the 3-fold axes, a channel-like structure at 5-fold axes and depressions at the icosahedral 2-fold axes. A 2/5-fold wall is located between the depression at the 2-fold axis and the 5-fold channel. (Figure 1a, b). More pronounced protrusions around 3-fold axis were observed in AAVPHP.eB capsid compared to those in AAV9 capsid (Figure 1b).

**Figure 1.**
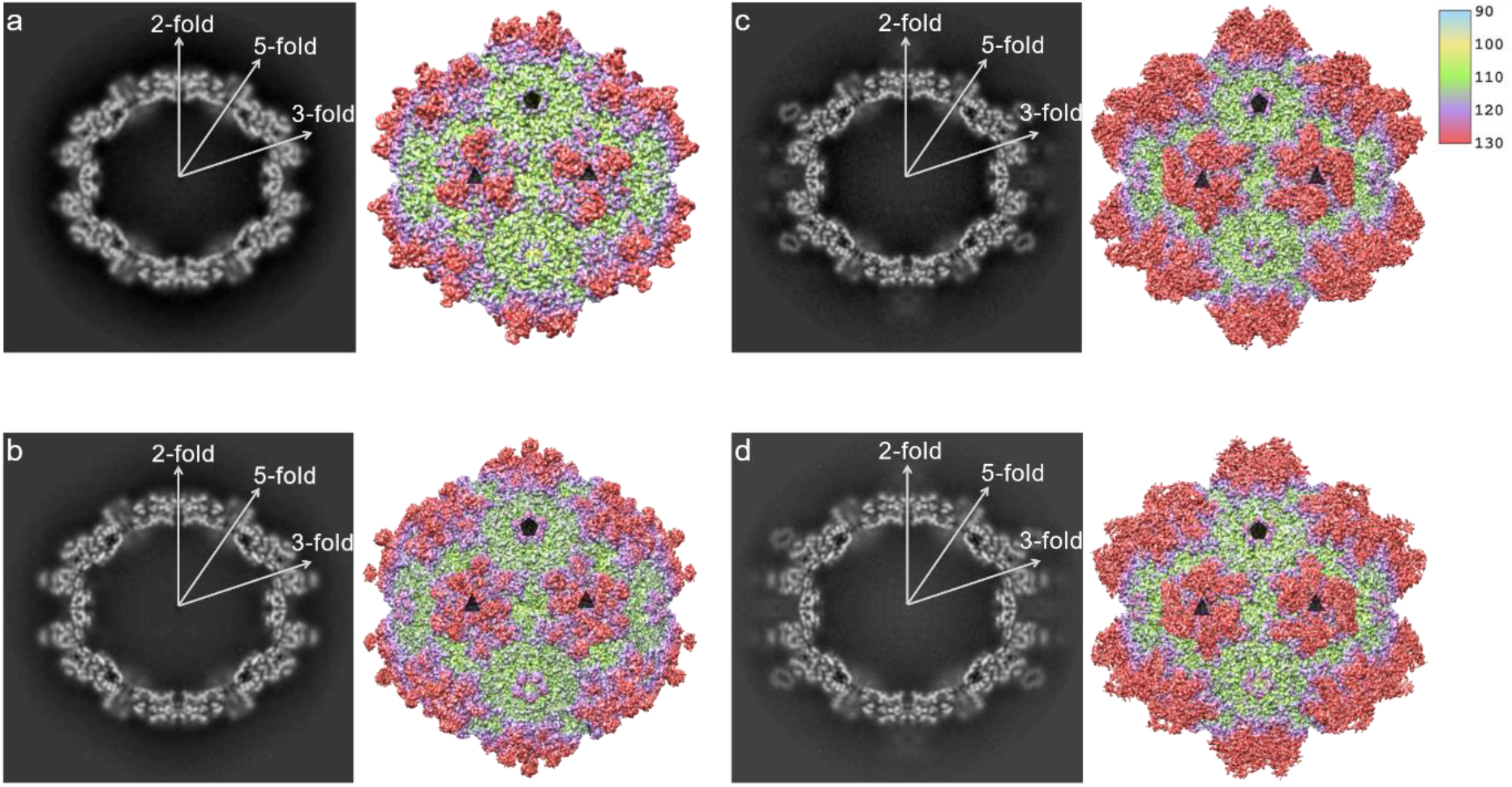
Cryo-EM reconstruction of AAV9 and AAVPHP.eB particles with or without AAVR binding. **(a)** AAV9, **(b)** AAVPHP.eB, **(c)** AAV9-AAVR and **(d)** AAVPHP.eB-AAVR. The central cross-sections are shown with the icosahedral two-, three- and fivefold axes. Density maps are radially colored by distance as shown in the color key and icosahedral 3-fold axis and 5-fold axis are represented by black triangles and pentagons.

Next, AAV9 and AAVPHP.eB were individually incubated with the soluble AAVR extracellular fragment containing PKD1-5 and structures of the AAV9-AAVR and AAVPHP.eB-AAVR complexes were subsequently determined by cryo-EM. The final resolutions of the cryo-EM reconstruction were estimated to be 3.23 Å for the AAV9-AAVR complex and 3.76 Å for the AAVPHP.eB-AAVR complex at a 0.143 cutoff of FSC **(Supplementary Table 1, Supplementary Figure 1)**. The resolution of additional attached density sitting above AAV9 and AAVPHP.eB capsid in AAVR complex reconstructions were sufficient to identified as AAVR PKD2 **(Supplementary Figure 2)**. Interaction pattern of AAVR PKD2 with AAV9 or AAVPHP.eB capsid is similar to that with AAV1 and AAV2^11,14^.

### AAV9 and AAVPHP.eB receptor binding interfaces

Resembling the engagement pattern of AAVR with AAV1 and AAV2, one PKD2 molecule also interacts with two capsid proteins of AAV9 or AAVPHP.eB. A total of 19 AAVR residues are within 4 Å distance of the AAV9 capsid and a total of 12 AAVR residues are within 4 Å distance of the AAVPHP.eB capsid **(Supplementary Table 2)**. Most AAV9 capsid interacting residues of PKD2 reside in A-B loop and B-C loop, three residues (R406, S413 and F416) in N-terminal of PKD2, two residues (I462 and K464) in D strand, and one additional residue (Y442) in C strand. While most AAVPHP.eB capsid interacting residues of PKD2 reside in the B-C loop, one residue (E418) in strand A, one residue (S425) in A-B loop and one residue (Y442) in C strand. Hydrogen-bond interacting residues prediction by LigPlot+ reveals that AAVR R406, S431, D435, D437 and I439 potentially form hydrogen bonds with AAV9 capsid, and a potential salt bridge between positively charged AAVR K438 and negatively charged AAV9 D384 (**Supplementary Table 2**, Figure 2a). Only four residues in AAVR PKD2 (S431, D435, D436, and D437) potentially form hydrogen bonds with the AAVPHP.eB capsid (Figure 2b). Surface plasmon resonance measurements indicate that AAV9 binds with AAVR at a KD of 138.5 nM in vitro, whereas KD of AAVPHP.eB is evaluated at 273.8nM **(Supplementary Figure 5)**. We reasoned that narrow interface, fewer hydrogen bonds and lack of salt bridge between AAVPHP.eB and AAVR PKD2 may provide molecular explanation for the lower affinity of AAVPHP.eB to AAVR.

**Figure 2.**
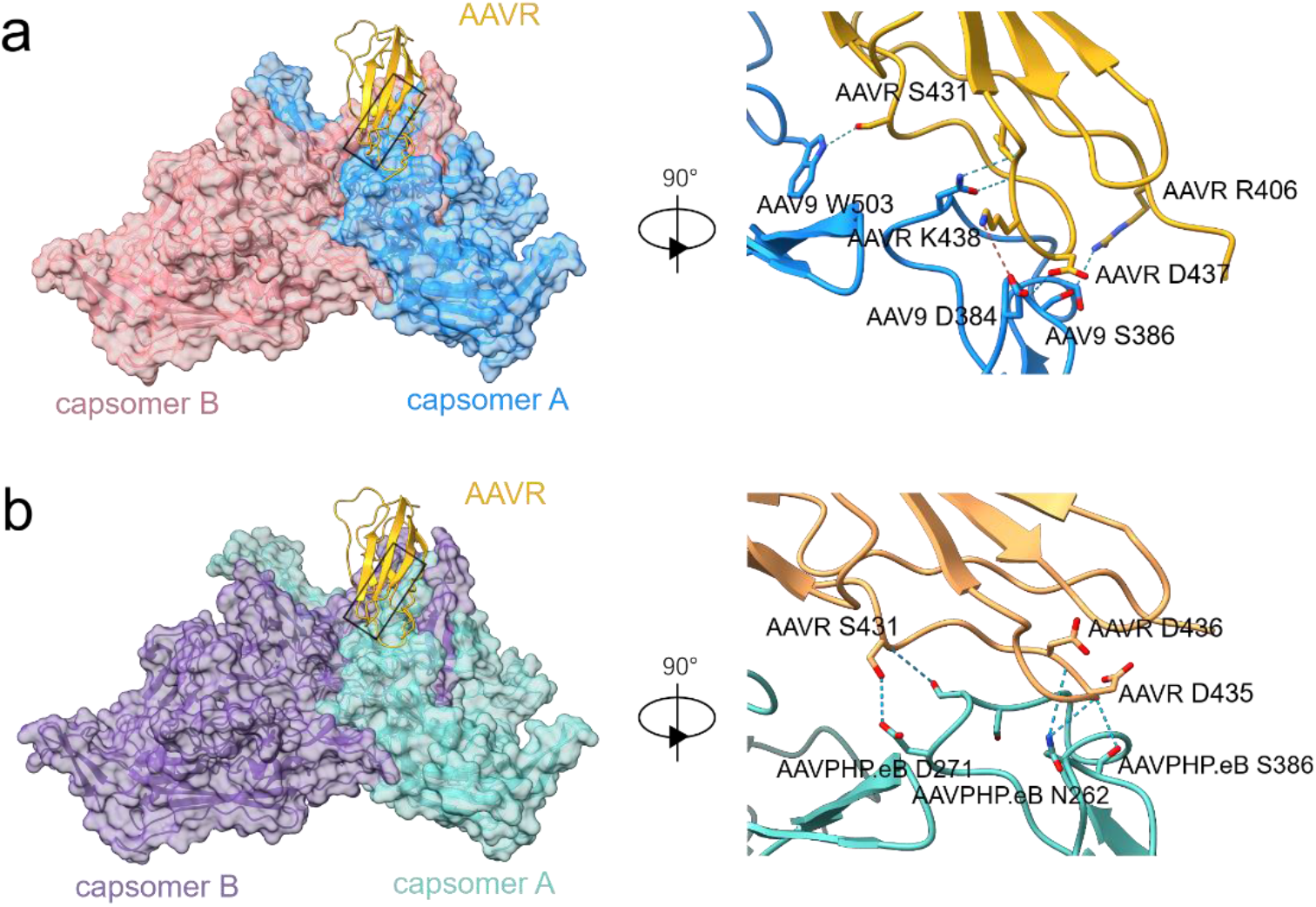
receptor interface of AAV9-AAVR and AAVPHP.eB-AAVR complexes. **(a)** One AAVR PKD2 (gold) interacts with two AAV9 capsomers (blue indication capsomer A, pink indicating capsomer B), and close-up view of the boxed region (90-degree rotation around Y axis). Only residues have potential side chain interaction are labeled in the diagram. **(b)**. One AAVR PKD2 (gold) interacts with two AAVPHP.eB capsomers (turquoise indicating capsomer A, purple indicating capsomer B), and close-up view of the boxed region (90-degree rotation around Y axis). Light blue dashes indicate potential hydrogen bonds, red dash indicates salt bridge.

### Capsid structure of AAV9 and AAVPHP.eB upon AAVR binding

Consistent with previously reported AAV structures, only viral protein 3 (VP3) common region density can be defined in AAV9 and AAVPHP.eB reconstruction maps. For AAV9, amino acid density from D219 to L736 was observable, and for AAVPHP.eB amino acid density from D219 to L743 was observable. In AAV9, there were 12 capsomer A residues and 4 capsomer B residues within 4 Å distance of AAVR PKD2. In AAVPHP.eB, 10 capsomer A residues and 3 capsomer B residues were within 4 Å distance of AAVR PKD2 **(Supplementary Table 2)**. All capsid residues close to AAVR PKD2 in both AAV9 and AAVPHP.eB reside in VR-I, VR-III, VR-IV, VR-V and VR-VIII.

The side-chain density of VR-VIII in AAV9 prior and post AAVR binding can be well defined at SD level of 1.5 (Figure 3a, b). While AAVPHP.eB possessed an engineered VR-VIII with an insertion of seven peptides (TLAVPFK) between the residue 588 and 589 and 2 mutations (A587D, Q588G AAVPHP.eB numbering)^7^. The 7-amino-acid insertion pointed further out from the capsid and does not alter the conformation of the ascending arm before S586 and descending arm after A596. The density of residues reside at the base of native AAVPHP.eB VR-VIII insertion was evident at SD level of 0.65, but L590 and A591 which reside on the top of the engineered loop remain lack of density under same SD level (Figure 3c). Upon AAVR binding, the main-chain density of L590 and A591 was revealed at SD level of 0.65 and the VR-VIII density can be better defined (Figure 3d). The density of other AAVR PKD2 interacting VRs (VR-I, VR-III, VR-IV and VR-V) in AAV9 and AAVPHP.eB native or AAVR bound state can be well defined at SD level of 1.

**Figure 3.**
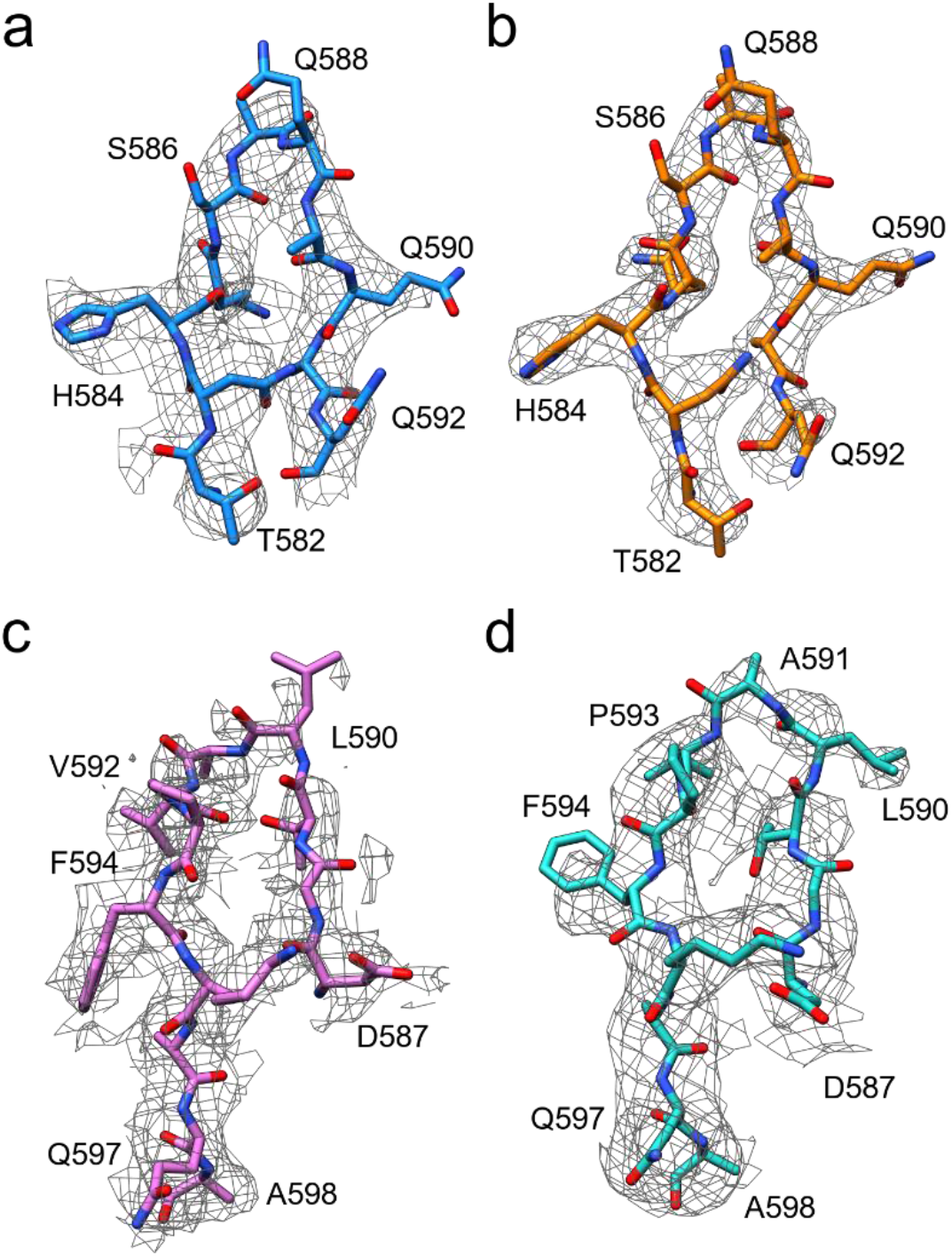
Density maps of VR-VIII. (a) AAV9, (b) AAV9-AAVR, (c) AAVPHP.eB and (d) AAVPHP.eB-AAVR VR-VIII models fitted into density maps. Electron density maps are shown in grey meshes.

Both AAV9 and AAVPHP.eB capsid protein show negligible overall conformational change upon AAVR binding. The capsid structures of native AAV9 and AAV9 complexed with AAVR shared a RMSD of 0.480 across all Cɑ atoms in 518 residues. And the capsid structures of AAVPHP.eB and AAVPHP.eB complexed with AAVR differed by a RMSD of 0.515 across all Cɑ atoms in 527 residues.

However, the difference at VR-I was much significant with a Cɑ RMSD of ~1.158 Å in AAV9 and ~1.152 Å in AAVPHP.eB. S268 main chain flipped away from AAVR and Cα moved away from receptor by ~2.95 Å in AAV9 and ~2.03 Å in AAVPHP.eB respectively upon AAVR binding (Figure 4 a, c). In the previous study, S268 in AAVrh.10 was proposed to be key a residue in BBB penetration^15,16^. Interestingly, S268 is also conserve in AAV9 and AAVPHP.eB sequence. The repulsion of S268 in AAV9/AAVPHP.eB upon AAVR binding and the sequence conversation of this residue among different BBB penetrating AAVs hint its special role in AAVR binding and the potential relationship between the AAVR recognizing and BBB penetrating of AAV9.

**Figure 4.**
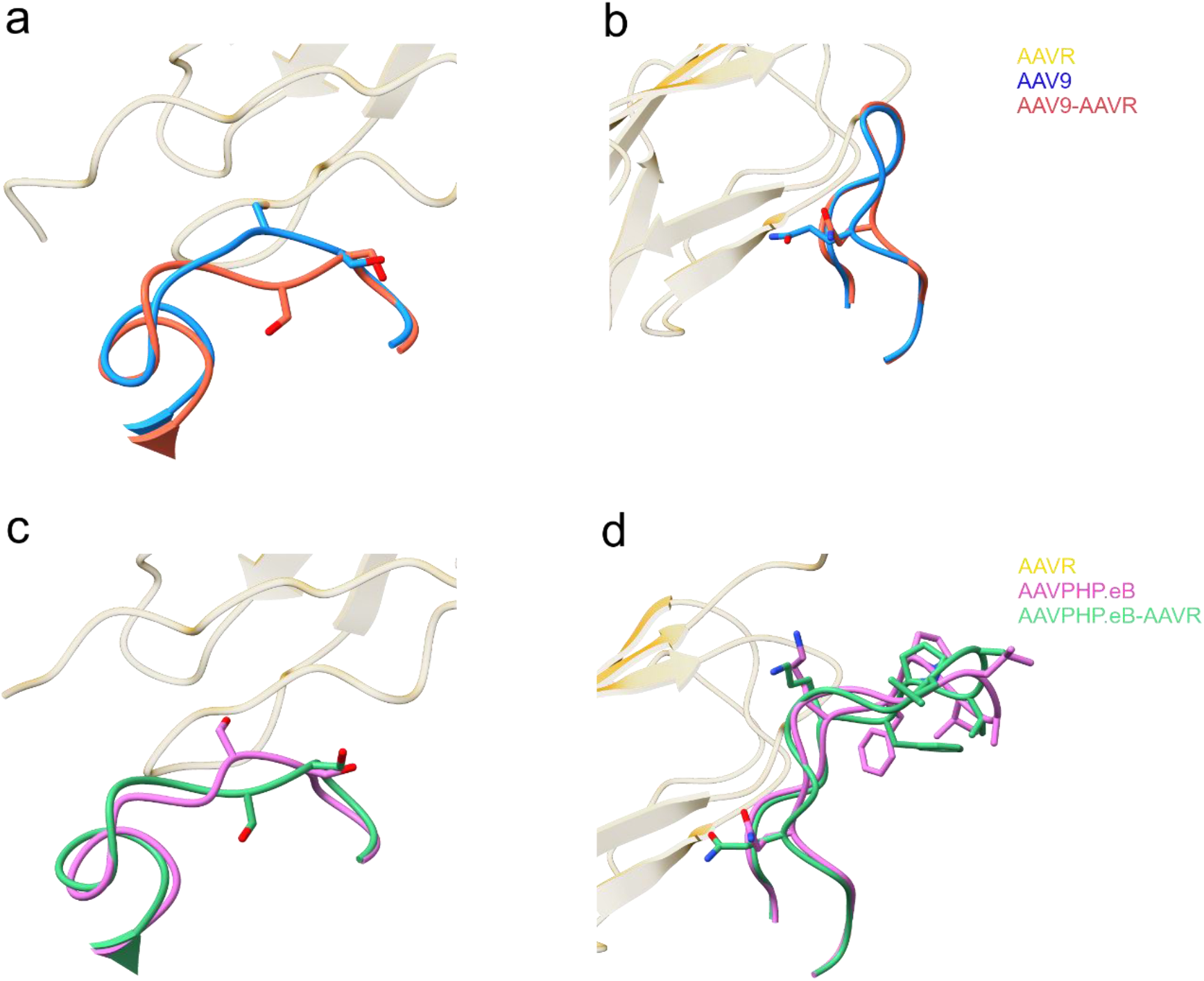
Conformational changes of AAV9/PHP.eB capsids upon AAVR binding. Conformational changes of (a)AAV9 and (c) AAVPHP.eB VR-I; VR-VIII of (b)AAV9 and (d) AAVPHP.eB are shown in ribbon representation. Side chains are shown in stick representation in same color as in main chain ribbon diagram.

The AAV9 VR-VIII (Cα RMSD ~0.76 Å) underwent less significant conformational change compared to AAVPHP.eB (Cα RMSD ~1.476 Å) AAVR binding. AAV9 Q585 was pushed away from AAVR by ~1.62 Å, while other residues in AAV9 VR-VIII showed no significant position shift (Figure 4 b). For AAVPHP.eB VR-VIII, P593 was pushed away from AAVR by ~1.53 Å and the apex for VR-VIII loop was lifted after AAVR interaction (Figure 4 d).

Collectively, these results suggest that the BBB penetrating associated reside S268 in VR-I may play special role in AAVR interaction, thus further indicate the potential relationship between AAVR and various tissue tropism.

### Diverse VR conformation in various AAV serotypes

To characterize similarities and differences between AAVR PKD2 interacting and BBB penetrating AAVs, we superposed AAV1, AAV2, AAV9, AAVPHP.eB and AAVrh.10 capsid structures. Superposition suggested that most structural variability occurred in VR-I, VR-II, VR-IV and VR-VIII.

AAVR PKD2 interacting VR-I and VR-IV showed most structural diversity among serotypes. Comparing with AAV2 VR-I, AAV1 and AAV9/PHP.eB VR-I exhibited more extended loop toward virus surface due to a single amino acid insertion in AAV1 VR-I and a 2-amino-acid insertion in AAV9/PHP.eB. Despite high sequence conservation among AAVrh.10 and AAV9/PHP.eB VR-I, AAVrh.10 VR-I exhibited a different conformation, which pointed outwards from the virus surface.

Among AAVR PKD2 interacting AAVs, only AAV9/PHP.eB have potential VR-IV interaction with AAVR. AAV9/PHP.eB VR-IV is closer to adjacent AAVR compared to that of AAV1. AAV2 VR-IV adopts a perpendicular position and more extended conformation compared to that of AAV1 and AAV9/PHP.eB. AAVrh.10 VR-IV has a similar loop conformation with that of AAV1.

VR-IIs located around 5-fold axis also exhibit structural discrepancy among serotypes. To note, although VR-II amino acids were identical in AAV9 and AAVPHP.eB, loops showed different conformation.

Despite the 7-amino-acid insertion and 2 mutation in VR-VIII of AAVPHP.eB, VR-VIIIs among different AAV serotypes shared common morphology.

### Relationship between AAV9 receptors and neutralizing antibody

Previous studies have identified galactose as primary attachment receptor for AAV9 and mapped N470, D271, N272, Y446, and W503 as the binding pocket at the region in between 3-fold protrusion and 2/5-fold wall^17, 18^. While the AAVR PKD2 footprint bridges the 3-fold protrusion and 2/5-fold wall on AAV9 capsid. The AAVR PKD2 footprint and galactose binding pocket only share one common residue W503, indicating galactose and AAVR may serve as independent receptor for AAV9 attachment and transduction.

To date, two antibody-AAV9 complex structures have been reported, including a BALB/c mouse originated and hybridoma-screening based antibody PAV9.1 and a commercially available nanobody CSAL9, both of which exhibit neutralizing activity against AAV9^19, 20^. The epitope of PAV9.1 lays on the 3-fold axis of AAV9 overlapping with AAVR footprint by two residues (Q588 and Q590) (Figure 5c). Another study powered by M13 phage display technology characterized potential immunogenic AAV9 VP3 epitopes largely overlap with the AAVR footprint regions on 3-fold protrusion and 2/5-fold wall^21^.

**Figure 5.**
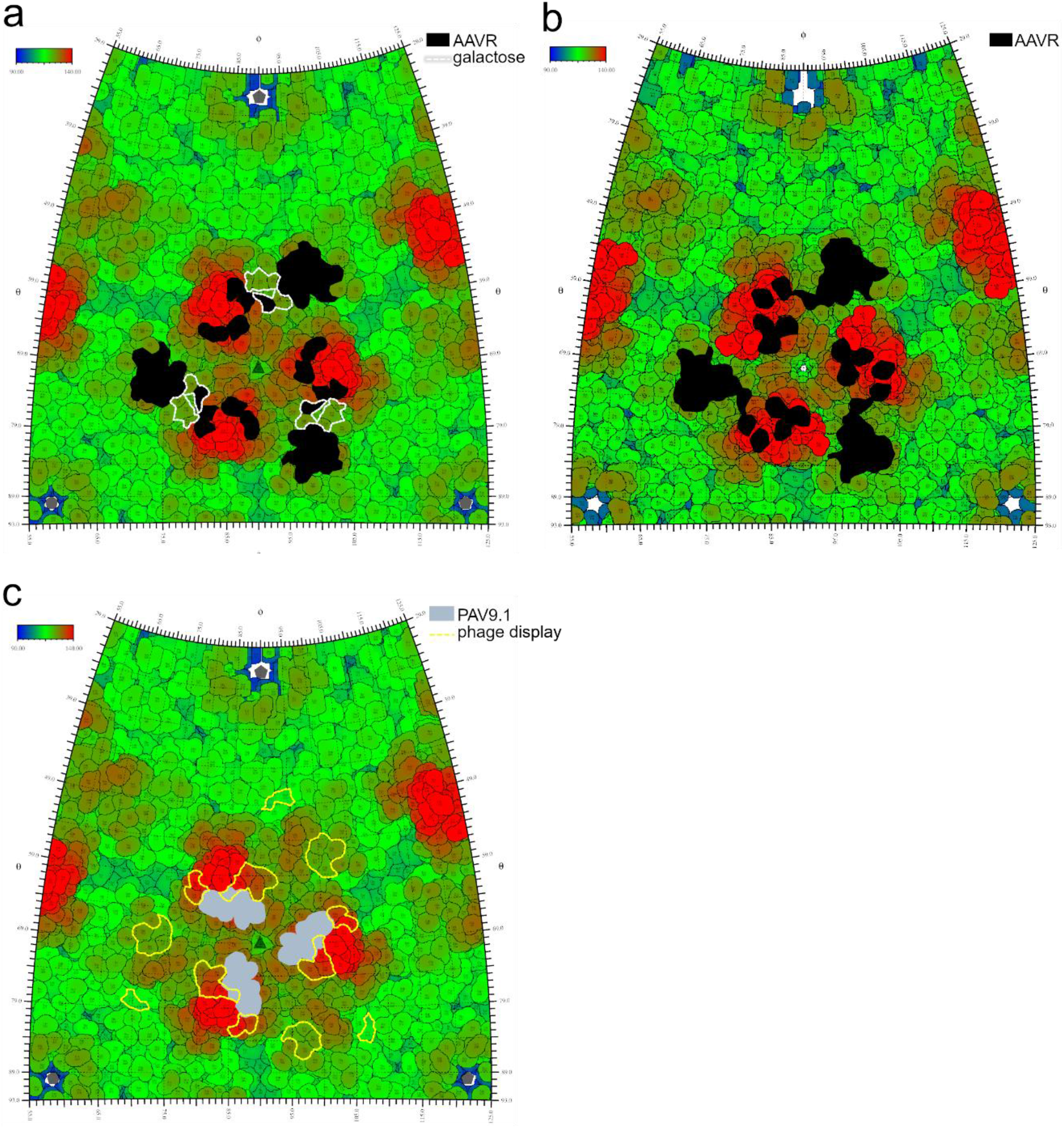
Receptor and antibody binding footprint on AAV9 and AAVPHP.eB capsid. Projections of the AAV9 and AAVPHP.eB surface. In (a) AAV9, AAVR PKD2 footprint is filled in black and galactose footprint is out lined in white. In (a) AAVPHP.eB, AAVR PKD2 footprint is filled in black. On (c) AAV9, PAV9.1 footprints are filled in grey and phage display screened immunogenetic peptides are outlined in yellow. Roadmaps was generated by RIVEM, the two angles (θ, φ) define a vector and a further location on the icosahedron surface. As show by the key, roadmaps are colored by distance from the center of the virus from blue (radius = 90Å) to red (radius = 140Å).

**Figure 6.**
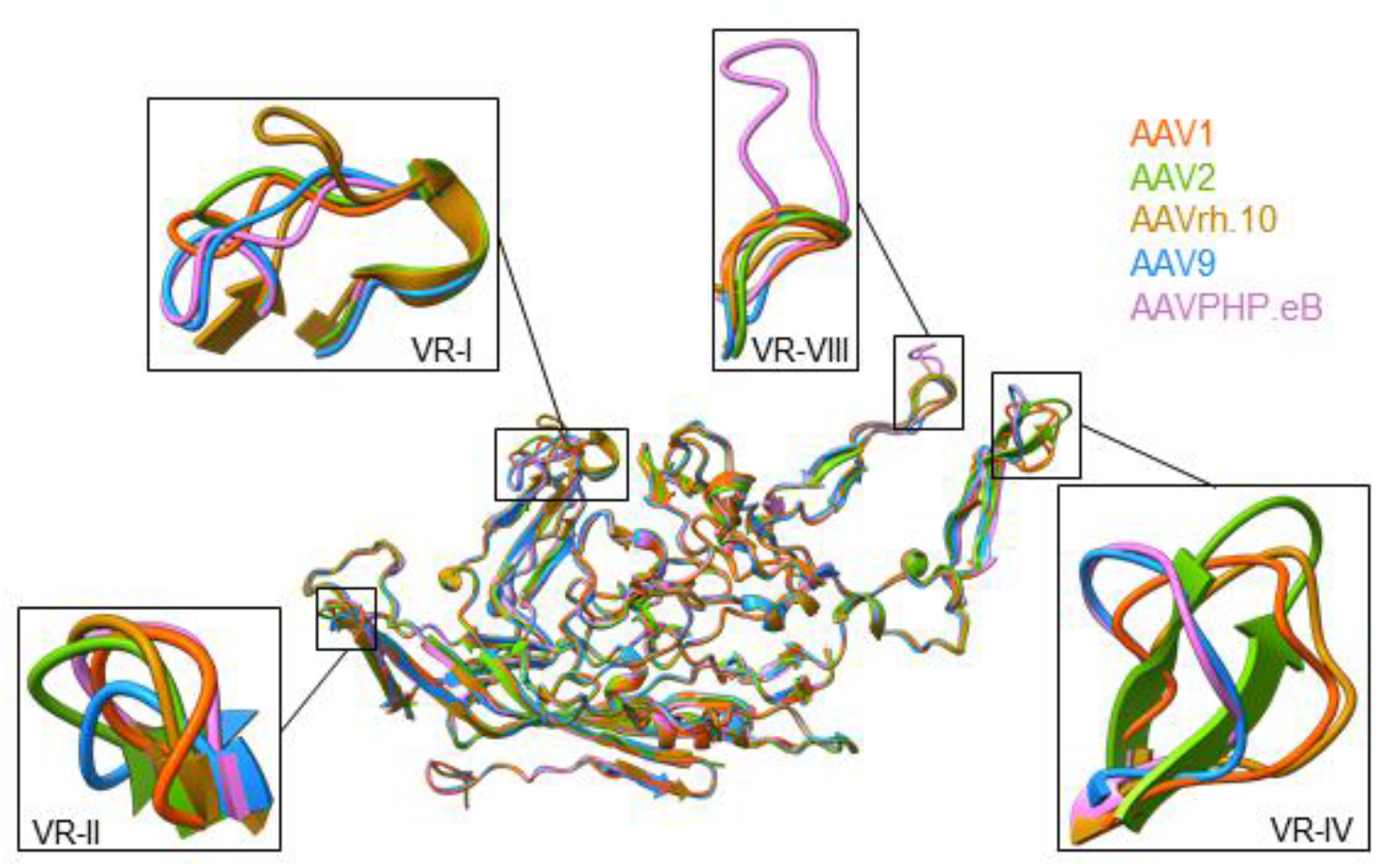
Structural superposition of VPs. Superposition of VP structures from AAV1, 2, 9, PHP.eB and rh.10. black boxes indicate the enlarged VR-I, VR-II, VR-IV and VR-VIII.

Interestingly, both AAVR PKD2 footprint and epitopes screened from phage display technology only overlap with galactose binding pocket by 1 residue (W503) on AAV9 capsid. Again, indicating W503 may serve as important dual functional residue in both galactose mediated viral attachment and AAVR mediated transduction. This is also in consistent with the finding in previous mutagenesis study which suggested a dual role of residues from E500 to W503 facilitating successful transduction other than galactose attachment^22^.

To note, CSAL9 binding to AAV9 also occludes residues around 5-fold axis and phage display screened epitopes also elicited antigenic VP1/2 N terminal residues^20, 21^, suggesting that other than AAVR, VP1/2 common N terminal region may also serve as crucial part in AAV transduction process.

### LY6A binding to AAVPHP.eB

Recent studies have reported that the enhanced CNS transduction and BBB penetration of AAVPHP.eB is driven by a GPI-anchored protein, LY6A, independent of AAVR. Then we purified extracellular domain of LY6A in 293F cells to further investigate their interaction. SPR sensorgrams reveal a small-molecule-like fast association and disassociation of LY6A to AAVPHP.eB **(Supplementary Figure 8b)**. Whereas injecting LY6A onto the AAV9 immobilized CM5 sensor chip, no SPR signals were detected **(Supplementary Figure 8a)**. Due to the dynamic interaction of LY6A to AAVPHP.eB, we incubated LY6A and AAVPHP.eB in the absence of AAVR with glutaraldehyde at a final concentration of 0.05% immediately before applying to cryo sample grids. Then the preliminary structure of AAVPHP.eB-LY6A is characterized by cryo-EM at 200kV. An additional vague density is observed on the top of AAVPHP.eB 3-fold axis and did not occupy the binding position of AAVR PKD2 **(Supplementary Figure 8c)**. To further validate LY6A binding site on AAVPHP.eB, a SPR competition assay was performed. Analysis of the resultant sensorgrams showed that the binding of AAVR had no impact on subsequent LY6A binding and vice versa **(Supplementary Figure 9)**. Additionally, this density extended from the middle of the 3-fold axis to interact with 3-fold protrusion formed by VR-VIII **(Supplementary Figure 8d and e)**. These observations further demonstrate that the enhanced CNS transduction and BBB penetration of AAVPHP.eB is driven by the alternative receptor LY6A independent of AAVR.

## Discussion

In this study, we report the native AAV9 and AAVPHP.eB and their complex structures with cellular receptor AAVR. The structures of AAV9-AAVR and AAVPHP.eB-AAVR demonstrate that AAVR PKD2 binds to AAV9/PHP.eB between 3-fold protrusion and 2/5-fold wall, which share a similar AAVR interaction pattern with that of AAV1 and AAV2.

VR-I, VR-III, VR-IV, VR-V and VR-VIII in AAV9 and AAVPHP.eB engaged with AAVR PKD2, which differ with AAVR PKD2 interacting VRs in AAV1 and AAV2 by VR-IV and VR-V. VR-IVs exhibit most conformational variance and AAV9/PHP.eB. VR-IVs are spatially closer in distance to adjacent AAVR to facilitate interaction. Despite common morphology among VR-Vs in different AAV serotypes, E500 and W503 in AAV9 VR-V have potential contacts with AAVR PKD2. To note, W503 also play a role in galactose binding in AAV9. Our structural finding further supports the notion that E500-W503 in the AAV9 capsid possess dual function for galactose binding and virus post-attachment process in previous study^22^.

The hybridoma screened PAV9.1 epitope footprint slightly overlaps with that of AAVR PKD2 on AAV9 (Q588 and Q590), and the immunogenetic peptides screened by phage display technique exhibit large overlap with AAVR PKD2 footprint on AAV9 VR-III and VR-V. This result indicates one antibody neutralizing mechanism is to occlude AAVR binding and virus cellular trafficking. However, another commercial antibody CASL9 epitopes reside around AAV9 5-fold axis, and the rest phage display screened immunogenetic peptides also suggest VP1 unique and VP1/2 common regions are antigenic, hint the alternative neutralizing mechanism through interfering the AAV cellular trafficking process other than AAVR binding.

AAV9 and its engineered variant AAVPHP.eB support the transduction of CNS and are able to cross BBB. Their AAVR bound complex structures revealed that S268 in VR-I underwent a conformational change upon AAVR binding. AAV9/PHP.eB S268 is also equivalent to S269 in AAVrh.10 which is reported to be important for BBB penetrating ability. However, despite the VR-I difference between AAV9 and AAVPHP.eB, little difference in AAVR interaction was observed including the engineered VR-VIII in AAVPHP.eB. Recent studies also further elucidate that AAVR is more likely to act as an entry factor participating in virus intracellular trafficking. In the light of studies on new acquired LY6A binding ability of AAVPHP.eB, we tried to further explore the molecular basis of AAVPHP.eB interaction with LY6A. We found that the engineered VR-VIII in AAVPHP.eB had potential interaction with LY6A. Suggesting an engineered VR-VIII facilitated alternative receptor recognition ability for AAVPHP.eB.

In summary, the structures of neurotropic clade F AAV9 and its engineered variant AAVPHP.eB in complexed with AAVR deepened the understanding of AAV receptor engagement and one dominant neutralizing mechanism. Structural analysis of AAV9-AAVR and AAVPHP.eB-AAVR suggest S268 in VR-I as determinant residue in BBB penetration. And structure information of AAVPHP.eB-LY6A also indicate that the enhanced CNS transducing character is facilitated by novel receptor recognition independent of AAVR. Our structure information would provide insights for vector engineering in attempts for higher transduction efficiency and altered tissue tropism.

## Methods

### Virus production and purification

Triple-plasmid transfection using polyethylenimine reagent (PEIMAX) (No. 24765, Polysciences, USA) was carried out to produce recombinant AAV9 and AAV-PHP.eB according to a previously reported procedure with modifications^11, 14^. Briefly, The plasmids pAAV9-GFP or pAAV-PHP.eB-GFP; pRepCap with AAV9 or AAV-PHP.eB encoding the Rep and Cap proteins; and pHelper plasmids were co-transfected into HEK293T cells. Cells were harvested 72 hr post-transfection, then AAV were purified with iodixanol gradient centrifugation. AAV genome copy titers were determined by real-time quantitative PCR (qPCR) using primers specific for the GFP gene sequences. The primers used were as follows: qpcr-GFP-F: TCTTCAAGTCCGCCATGCC; qpcr-GFP-R: TGTCGCCCTCGAACTTCAC.

### Purification of AAVR proteins

cDNAs encoding the AAVR PKD1-5 domains with a C-terminal His-tag in a pET28a vector were transformed into Escherichia coli BL21 (DE3) cells harboring the recombinant plasmids were cultured in Luria-Bertani (LB) medium containing 50 μg/ml kanamycin at 37 °C. Protein expression was induced by the addition of isopropyl 0.5 mM β-D-thiogalactoside (IPTG) at the OD600 of 0.6, followed by another 16-hr of cell culture. Protein purification was performed according to the previous reports^11, 14^. Briefly, recombinant protein was initially purified by nickel affinity chromatography (Qiagen, Holland) and subsequent size exclusion using Superdex 200 increase (GE Healthcare, USA), collected the peak around 14 ml. The purified proteins were concentrated to 6 mg/ml for storage at −80 °C until use.

### Sample preparation and cryo-EM data collection

AAV9 or AAV-PHP.eB particles and purified wt AAVR were mixed at a molar ratio of 1:120 (AAV:AAVR) at 4 °C for 1hr. An aliquot of 3 μl of each mixture was loaded onto a glow-discharged, carbon-coated copper grid (GIG, Au 2/1 200 mesh; Lantuo, China) bearing an ultrathin layer of carbon. The grid was then blotted for 4.5 s with a blot force of 0 in 100% relative humidity and plunge-frozen in liquid ethane using a Vitrobot Mark IV (FEI, USA). Cryo-EM data were collected with a 200 kV Arctica D683 electron microscope (FEI, USA) and a Falcon II direct electron detector (FEI, USA). A series of micrographs were collected as movies (19 frames, 1.2 s) and recorded with −2.2 to −0.5 μm defocus at a calibrated magnification of 110,000×, resulting in a pixel size of 0.93 Å per pixel. Statistics for data collection and refinement are summarized in Supplementary Table 1.

### Image processing and three-dimensional reconstruction

Similar image processing procedures were employed for all data sets. Individual frames from each micrograph movie were aligned and averaged using MotionCor2^23^ to produce drift-corrected images. Particles were picked and selected in RELION 2.1^24^, and the contrast transfer function (CTF) parameters were estimated using CTFFIND4^25^. Subsequent steps for particle picking and 2D and 3D classification were performed with RELION 2.1. The final selected particles (for AAVR complex samples, particles with clear additional densities above viral particles were selected after 3D classification) were reconstructed with THUNDER^26^. For all reconstructions, the final resolution was assessed using the gold-standard FSC criterion (FSC = 0.143) with RELION 2.1.

### Model building and refinement

To solve the structure of AAV9 and AAV-PHP.eB, the X-ray crystal structure of AAV9 (PDB code: 3ux1)^27^ was manually placed and rigid body fitted into the cryo-EM density map with UCSF Chimera^28^. To solve the AAV9-AAVR and AAVPHP.eB-AAVR complexes, the PKD2 domain structure from the AAV2-AAVR structure (PDB: 6IHB) was manually aligned with cryo-EM density corresponding to the bound receptors. Manual adjustment of amino acids of AAV9/PHP.eB and PKD2 was performed using Coot^29^ in combination with real space refinement with Phenix. The data validation statistics shown in **Supplementary Table 1** were reported by MolProbity using the integrated function within the Phenix statistics module^30^.

### Surface plasmon resonance (SPR)

SPR analyses were carried out using a Biacore T200 (GE Healthcare, USA) with a flow rate of 30 μl/min at 25 °C in PBS buffer. AAV9 or AAVPHP.eB particles suspended in sodium acetate buffer (pH 4.0) were immobilized on a CM5 sensor chip by amide coupling. Different concentrations of the recombinant wildtype AAVR protein flowed over the chip and between each sample we used Glycine-HCl (10mM Glycine, pH2.0) for chip surface regeneration. The binding affinity was determined by and curves were generated by BIAEvaluation software (GE Healthcare, USA).

### SPR binding competition assay

Binding competition assays were performed by SPR (Biacore S200, GE). A-B-A injection method were used to unravel if LY6A and AAVR will simultaneously bind to AAVPHP.eB capsid. The AAVPHP.eB capsid was immobilized as described above. The A-B-A was used with 90 s injections of analyte A to ensure saturation or near-saturation was reached prior to injection of analyte B. Then analyte B is injected with saturated concentration of first analyte A.

## Data availability

The cryo-EM density maps and the structures were deposited into the Electron Microscopy Data Bank (EMDB) and Protein Data Bank (PDB) with the following accession numbers: AAV9 alone, XXXX; AAV9-AAVR, XXXX; AAVPGP.eB alone, XXXX; AAVPHP.eB-AAVR, XXXX. All other data supporting the findings of this study are available from the corresponding authors upon request.

## Correspondence

Correspondence and requests for materials should be addressed to G.X and Z.L.

## Acknowledgments

We thank the Computing and cryo-EM Platforms of Tsinghua University, Branch of the National Center for Protein Sciences (Beijing) for providing facilities. We thank Dr. Jianlin Lei and Mr. Tao Liu for their help in data collection. This work was supported by the National Program on Key Research Project of China (2020YFA0707500 and 2017YFC0840300), the National Natural Science Foundation of China (grants no. 31971126 and U20A20135), the China Postdoctoral Science Foundation (Grant BX2021165) and the Shuimu Tsinghua Scholar Program of Tsinghua University (Grant 2020SM142).

## Author contributions

Z.L. and G.X. conceived the project. Z.L. designed the experiments. R.Z., G.X., K.Y. and X.M performed experiments. R.Z., G.X., and Z.L. analyzed the data. Z.L. and G.X. wrote the manuscript. All authors discussed the experiments, read and approved the manuscript.

## References

1. Melchiorri, D., Pani, L., Gasparini, P., Cossu, G., Ancans, J., Borg, J.J., Drai, C., Fiedor, P., Flory, E., Hudson, I., et al. (2013). Regulatory evaluation of Glybera in Europe — two committees, one mission. Nat Rev Drug Discov 12, 719–719.

2. Russell, S., Bennett, J., Wellman, J.A., Chung, D.C., Yu, Z.-F., Tillman, A., Wittes, J., Pappas, J., Elci, O., McCague, S., et al. (2017). Efficacy and safety of voretigene neparvovec (AAV2-hRPE65v2) in patients with RPE65 -mediated inherited retinal dystrophy: a randomised, controlled, open-label, phase 3 trial. The Lancet 390, 849–860.

3. Al-Zaidy, S., Pickard, A.S., Kotha, K., Alfano, L.N., Lowes, L., Paul, G., Church, K., Lehman, K., Sproule, D.M., Dabbous, O., et al. (2019). Health outcomes in spinal muscular atrophy type 1 following AVXS-101 gene replacement therapy. Pediatr Pulmonol 54, 179–185.

4. Wu, Z., Asokan, A., and Samulski, R.J. (2006). Adeno-associated Virus Serotypes: Vector Toolkit for Human Gene Therapy. Molecular Therapy 14, 316–327.

5. Terstappen, G.C., Meyer, A.H., Bell, R.D., and Zhang, W. (2021). Strategies for delivering therapeutics across the blood–brain barrier. Nat Rev Drug Discov.

6. Deverman, B.E., Pravdo, P.L., Simpson, B.P., Kumar, S.R., Chan, K.Y., Banerjee, A., Wu, W.-L., Yang, B., Huber, N., Pasca, S.P., et al. (2016). Cre-dependent selection yields AAV variants for widespread gene transfer to the adult brain. Nature Biotechnology 34, 204–209.

7. Chan, K.Y., Jang, M.J., Yoo, B.B., Greenbaum, A., Ravi, N., Wu, W.-L., Sánchez-Guardado, L., Lois, C., Mazmanian, S.K., Deverman, B.E., et al. (2017). Engineered AAVs for efficient noninvasive gene delivery to the central and peripheral nervous systems. Nature Neuroscience 20, 1172–1179.

8. Ravindra Kumar, S., Miles, T.F., Chen, X., Brown, D., Dobreva, T., Huang, Q., Ding, X., Luo, Y., Einarsson, P.H., Greenbaum, A., et al. (2020). Multiplexed Cre-dependent selection yields systemic AAVs for targeting distinct brain cell types. Nat Methods 17, 541–550.

9. Pillay, S., Meyer, N.L., Puschnik, A.S., Davulcu, O., Diep, J., Ishikawa, Y., Jae, L.T., Wosen, J.E., Nagamine, C.M., Chapman, M.S., et al. (2016). An essential receptor for adeno-associated virus infection. Nature 530, 108–112.

10. Pillay, S., Zou, W., Cheng, F., Puschnik, A.S., Meyer, N.L., Ganaie, S.S., Deng, X., Wosen, J.E., Davulcu, O., Yan, Z., et al. (2017). Adeno-associated Virus (AAV) Serotypes Have Distinctive Interactions with Domains of the Cellular AAV Receptor. Journal of Virology 91, e00391-17, /jvi/91/18/e00391-17.atom.

11. Zhang, R., Xu, G., Cao, L., Sun, Z., He, Y., Cui, M., Sun, Y., Li, S., Li, H., Qin, L., et al. (2019). Divergent engagements between adeno-associated viruses with their cellular receptor AAVR. Nat Commun 10, 3760.

12. Huang, Q., Chan, K.Y., Tobey, I.G., Chan, Y.A., Poterba, T., Boutros, C.L., Balazs, A.B., Daneman, R., Bloom, J.M., Seed, C., et al. (2019). Delivering genes across the blood-brain barrier: LY6A, a novel cellular receptor for AAV-PHP.B capsids. PLoS ONE 14, e0225206.

13. Hordeaux, J., Yuan, Y., Clark, P.M., Wang, Q., Martino, R.A., Sims, J.J., Bell, P., Raymond, A., Stanford, W.L., and Wilson, J.M. (2019). The GPI-Linked Protein LY6A Drives AAV-PHP.B Transport across the Blood-Brain Barrier. Molecular Therapy 27, 912–921.

14. Zhang, R., Cao, L., Cui, M., Sun, Z., Hu, M., Zhang, R., Stuart, W., Zhao, X., Yang, Z., Li, X., et al. (2019). Adeno-associated virus 2 bound to its cellular receptor AAVR. Nat Microbiol 4, 675–682.

15. Albright, B.H., Storey, C.M., Murlidharan, G., Castellanos Rivera, R.M., Berry, G.E., Madigan, V.J., and Asokan, A. (2018). Mapping the Structural Determinants Required for AAVrh.10 Transport across the Blood-Brain Barrier. Molecular Therapy 26, 510–523.

16. Mietzsch, M., Barnes, C., Hull, J.A., Chipman, P., Xie, J., Bhattacharya, N., Sousa, D., McKenna, R., Gao, G., and Agbandje-McKenna, M. (2019). Comparative Analysis of the Capsid Structures of AAVrh.10, AAVrh.39, and AAV8. J Virol 94, e01769-19, /jvi/94/6/JVI.01769-19.atom.

17. Shen, S., Bryant, K.D., Brown, S.M., Randell, S.H., and Asokan, A. (2011). Terminal N-Linked Galactose Is the Primary Receptor for Adeno-associated Virus 9*. Journal of Biological Chemistry 286, 13532–13540.

18. Bell, C.L., Gurda, B.L., Van Vliet, K., Agbandje-McKenna, M., and Wilson, J.M. (2012). Identification of the Galactose Binding Domain of the Adeno-Associated Virus Serotype 9 Capsid. Journal of Virology 86, 7326–7333.

19. Giles, A.R., Govindasamy, L., Somanathan, S., and Wilson, J.M. (2018). Mapping an Adeno-associated Virus 9-Specific Neutralizing Epitope To Develop Next-Generation Gene Delivery Vectors. J Virol 92, e01011-18, /jvi/92/20/e01011-18.atom.

20. Mietzsch, M., Smith, J.K., Yu, J.C., Banala, V., Emmanuel, S.N., Jose, A., Chipman, P., Bhattacharya, N., McKenna, R., and Agbandje-McKenna, M. (2020). Characterization of AAV-Specific Affinity Ligands: Consequences for Vector Purification and Development Strategies. Mol Ther Methods Clin Dev 19, 362–373.

21. Chew, W.L., Tabebordbar, M., Cheng, J.K.W., Mali, P., Wu, E.Y., Ng, A.H.M., Zhu, K., Wagers, A.J., and Church, G.M. (2016). A multifunctional AAV-CRISPR-Cas9 and its host response. Nat Methods 13, 868–874.

22. Adachi, K., Enoki, T., Kawano, Y., Veraz, M., and Nakai, H. (2014). Drawing a high-resolution functional map of adeno-associated virus capsid by massively parallel sequencing. Nat Commun 5, 3075.

23. Zheng, S.Q., Palovcak, E., Armache, J.-P., Verba, K.A., Cheng, Y., and Agard, D.A. (2017). MotionCor2: anisotropic correction of beam-induced motion for improved cryo-electron microscopy. Nat Methods 14, 331–332.

24. Scheres, S.H.W. (2012). RELION: Implementation of a Bayesian approach to cryo-EM structure determination. Journal of Structural Biology 180, 519–530.

25. Rohou, A., and Grigorieff, N. (2015). CTFFIND4: Fast and accurate defocus estimation from electron micrographs. Journal of Structural Biology 192, 216–221.

26. Hu, M., Yu, H., Gu, K., Wang, Z., Ruan, H., Wang, K., Ren, S., Li, B., Gan, L., Xu, S., et al. (2018). A particle-filter framework for robust cryo-EM 3D reconstruction. Nat Methods 15, 1083–1089.

27. DiMattia, M.A., Nam, H.-J., Van Vliet, K., Mitchell, M., Bennett, A., Gurda, B.L., McKenna, R., Olson, N.H., Sinkovits, R.S., Potter, M., et al. (2012). Structural Insight into the Unique Properties of Adeno-Associated Virus Serotype 9. J Virol 86, 6947–6958.

28. Pettersen, E.F., Goddard, T.D., Huang, C.C., Couch, G.S., Greenblatt, D.M., Meng, E.C., and Ferrin, T.E. (2004). UCSF Chimera--A visualization system for exploratory research and analysis. J. Comput. Chem. 25, 1605–1612.

29. Emsley, P., Lohkamp, B., Scott, W.G., and Cowtan, K. (2010). Features and development of Coot. Acta Crystallogr D Biol Crystallogr 66, 486–501.

30. Afonine, P.V., Klaholz, B.P., Moriarty, N.W., Poon, B.K., Sobolev, O.V., Terwilliger, T.C., Adams, P.D., and Urzhumtsev, A. (2018). New tools for the analysis and validation of cryo-EM maps and atomic models. Acta Crystallogr D Struct Biol 74, 814–840.

