## Supplementary material for "Structural basis for the neurotropic AAV9 and the engineered AAVPHP.eB recognition with cellular receptors": Suplemental materials

**Contents** **Page**

### Supplementary Figures

#### Supplementary Figure 1

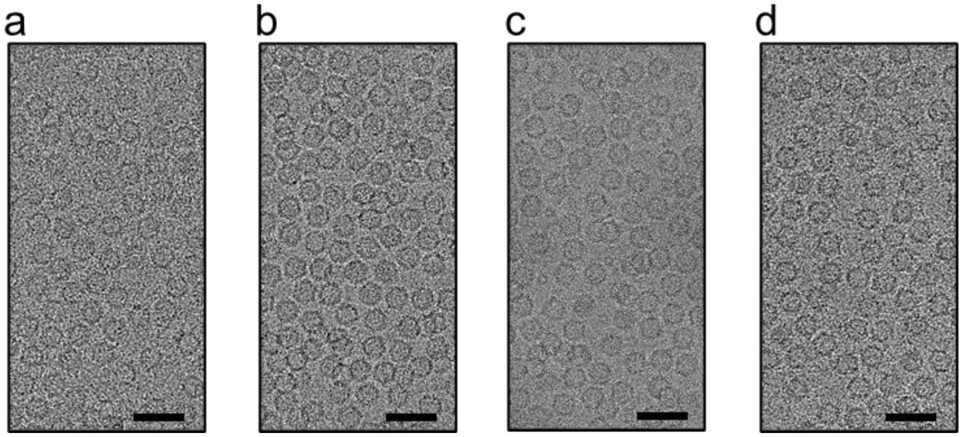

**Supplementary Figure 1.** Representative micrographs of (a) AAV9, (b) AAV9-AAVR, (c) AAVPHP.eB and (d) AAVPHP.eB-AAVR. Shown micrograph's size is 2048×4096 pixels cutted out from 4096x4096-pixels original micrographs. Scale bars indicate 50 nm.

**Supplementary Figure 2**

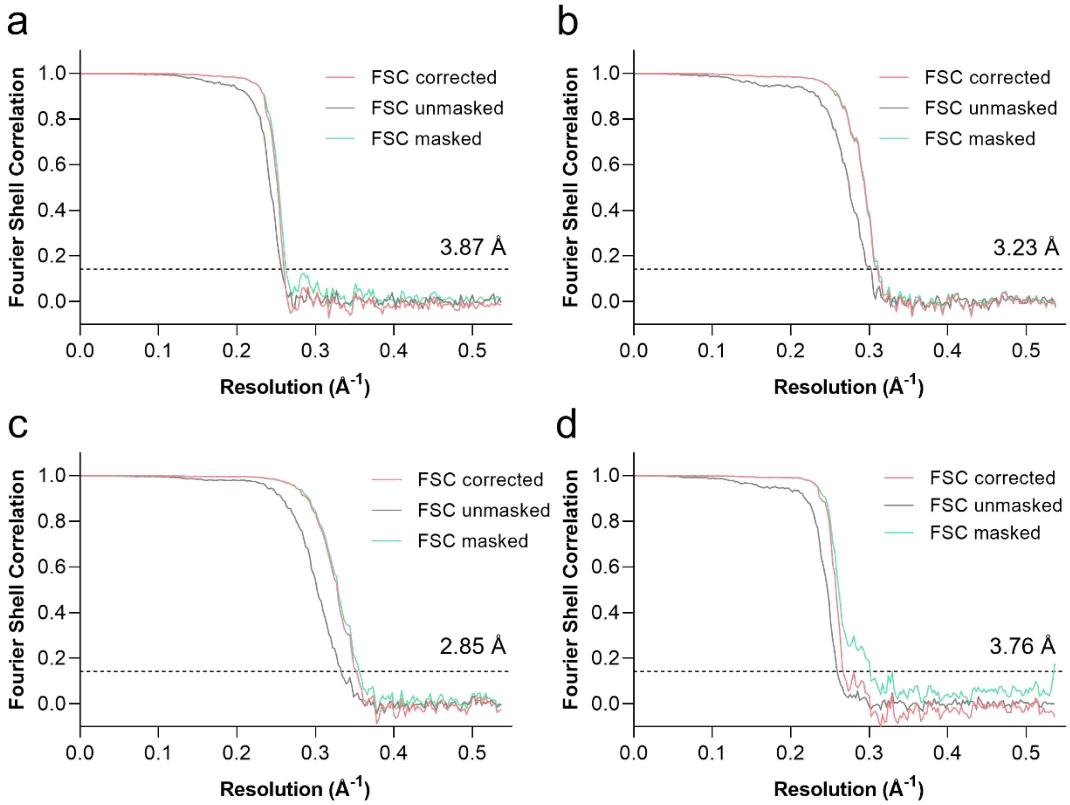

**Supplementary Figure 2. Resolution assessment.** Fourier shell correlation (FSC) of the final 3D reconstruction following gold standard refinement using RELION and THUNDER. The resolutions corresponding to an FSC of 0.143 are shown for unbound AAV9 **(a)**, the AAV9-AAVR complex **(b)**, unbound AAV-PHP.eB **(c)** and the AAV-PHP.eB-AAVR complex **(d)**. Unmasked FSC curves are colored in grey and masked FSC curves are colored in green. Corerected maps accounting for the effect of masking using phase randomization are colored in red.

**Supplementary Figure 3**

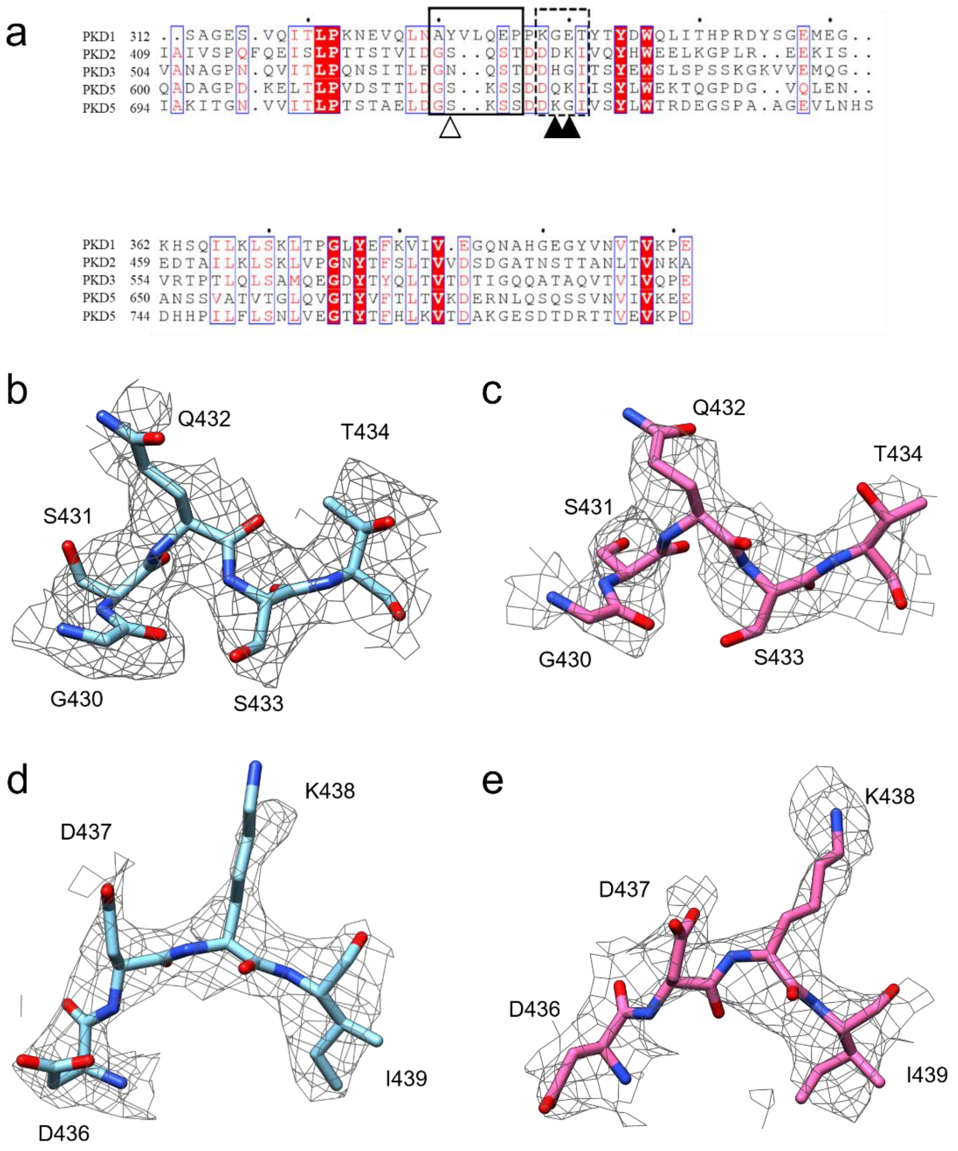

**Supplementary Figure 3.** (a)Sequence alignment of AAVR PKD1-5. Residues with red character or red backgrounds are conserved or identical, respectively. Signature resides are labeled with white and black triangles. Residues in AAVR PKD2 fitted in density in (b)(c) and (d)(e) are boxed in black line and black dash line, respectively. (a)AAV9-AAVR and (c)AAVPHP.eB-AAVR AAVR PKD2 feature fragments (G430~T434) are shown in stick diagrams and fitted into density represented by grey mesh. (a)AAV9-AAVR and (c)AAVPHP.eB-AAVR

AAVR PKD2 feature fragments (D436~I439) are shown in stick diagrams and fitted into density represented by grey mesh

**Supplementary Figure 4**

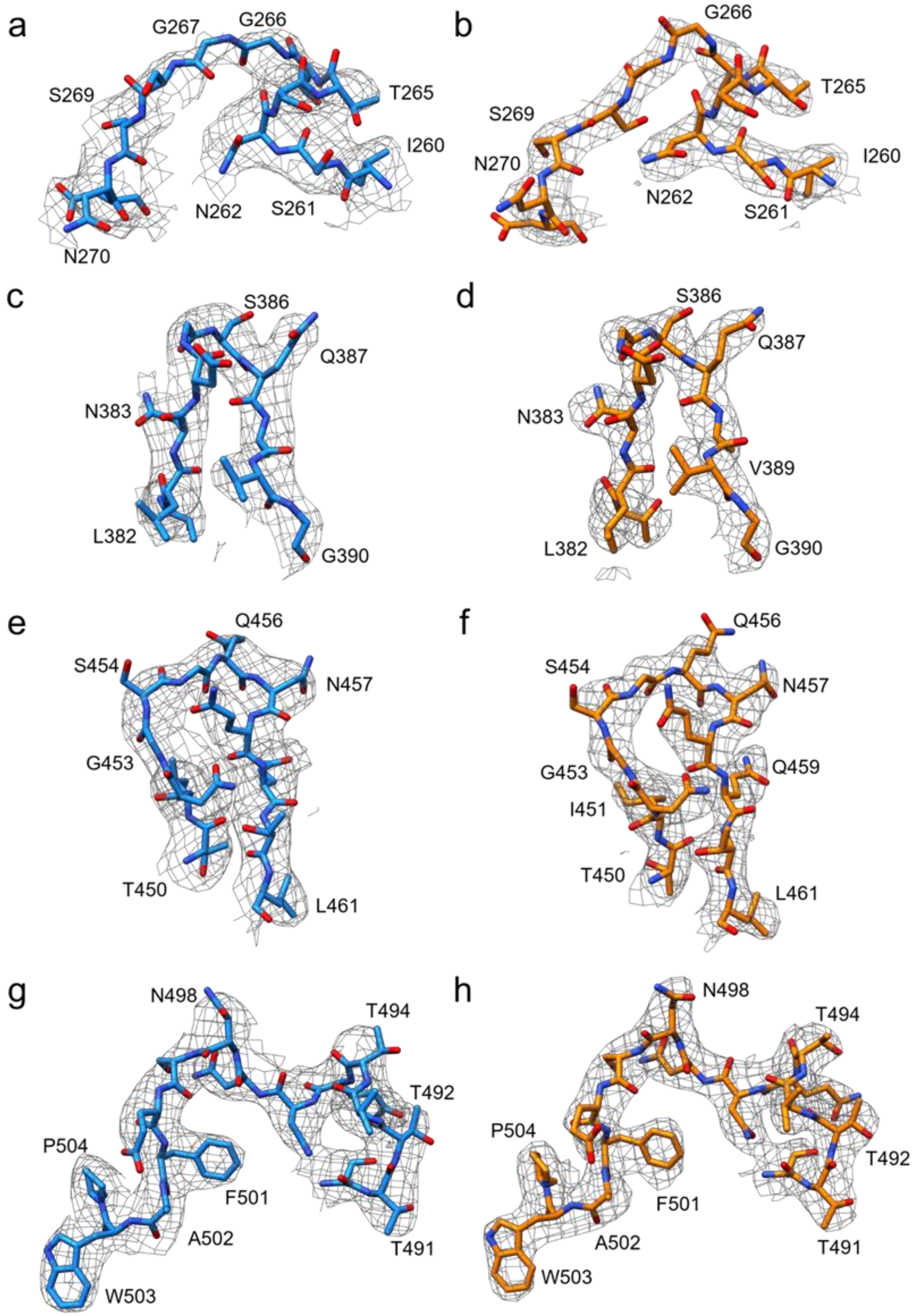

**Supplementary Figure 4.** Left side of the panel shows the native AAV9 (a)VR-I, (c)VR-III, (e)VR-IV and (g) VR-V fragments' stick models fitted into electron density. Right side of the panel shows the AAV9-AAVR complex (a)VR-I, (c)VR-III, (e)VR-IV and (g) VR-V fragments' stick models fitted into electron density. Electron density is presented as grey mesh.

**Supplementary Figure 5**

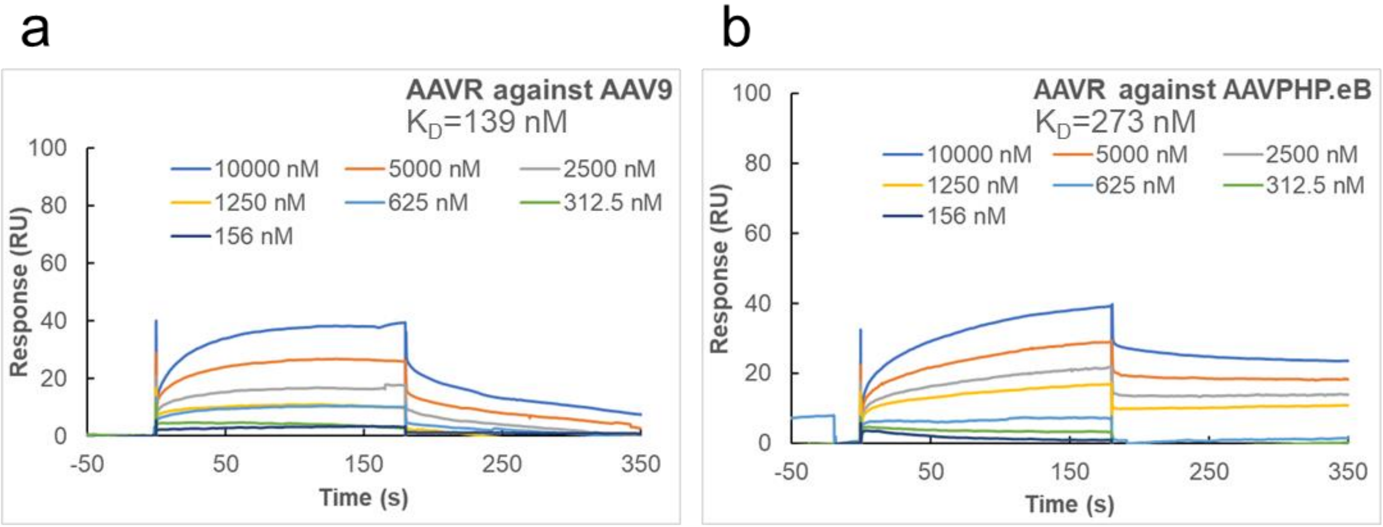

**Supplementary Figure 5.** SPR sensorgrams for binding of AAVR PKD1-5 as analyte to immobilized (a) AAV9 and (b) AAVPHP.eB on CM5 chip.

**Supplementary Figure 6**

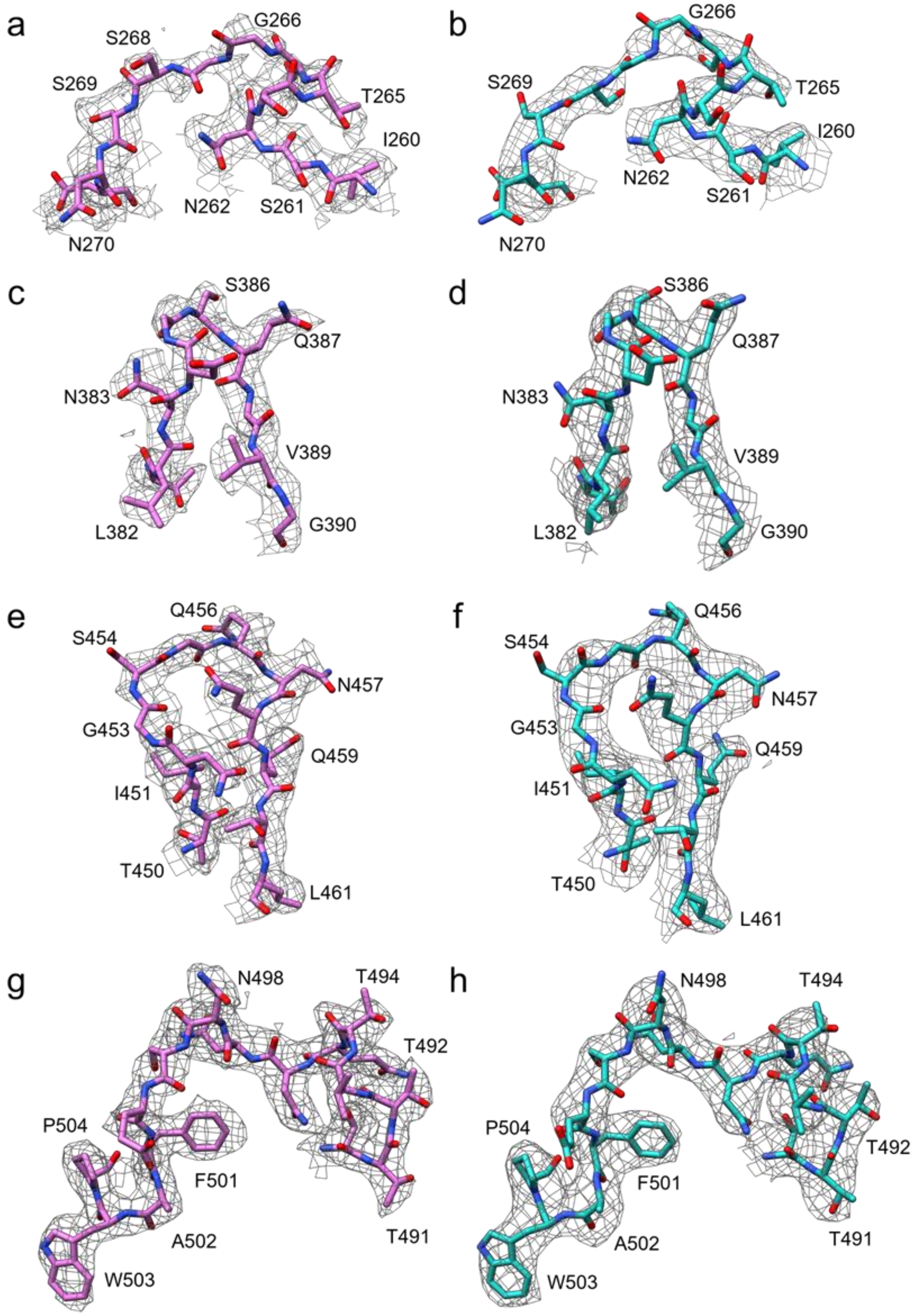

**Supplementary Figure 6.** Left side of the panel shows the native AAVPHP.eB (a)VR-I, (c)VR-III, (e)VR-IV and (g) VR-V fragments' stick models fitted into electron density. Right side of the panel shows the AAVPHP.eB-AAVR complex (a)VR-I, (c)VR-III, (e)VR-IV and (g) VR-V fragments' stick models fitted into electron density. Electron density is presented as grey mesh.

### Supplementary Figure 7

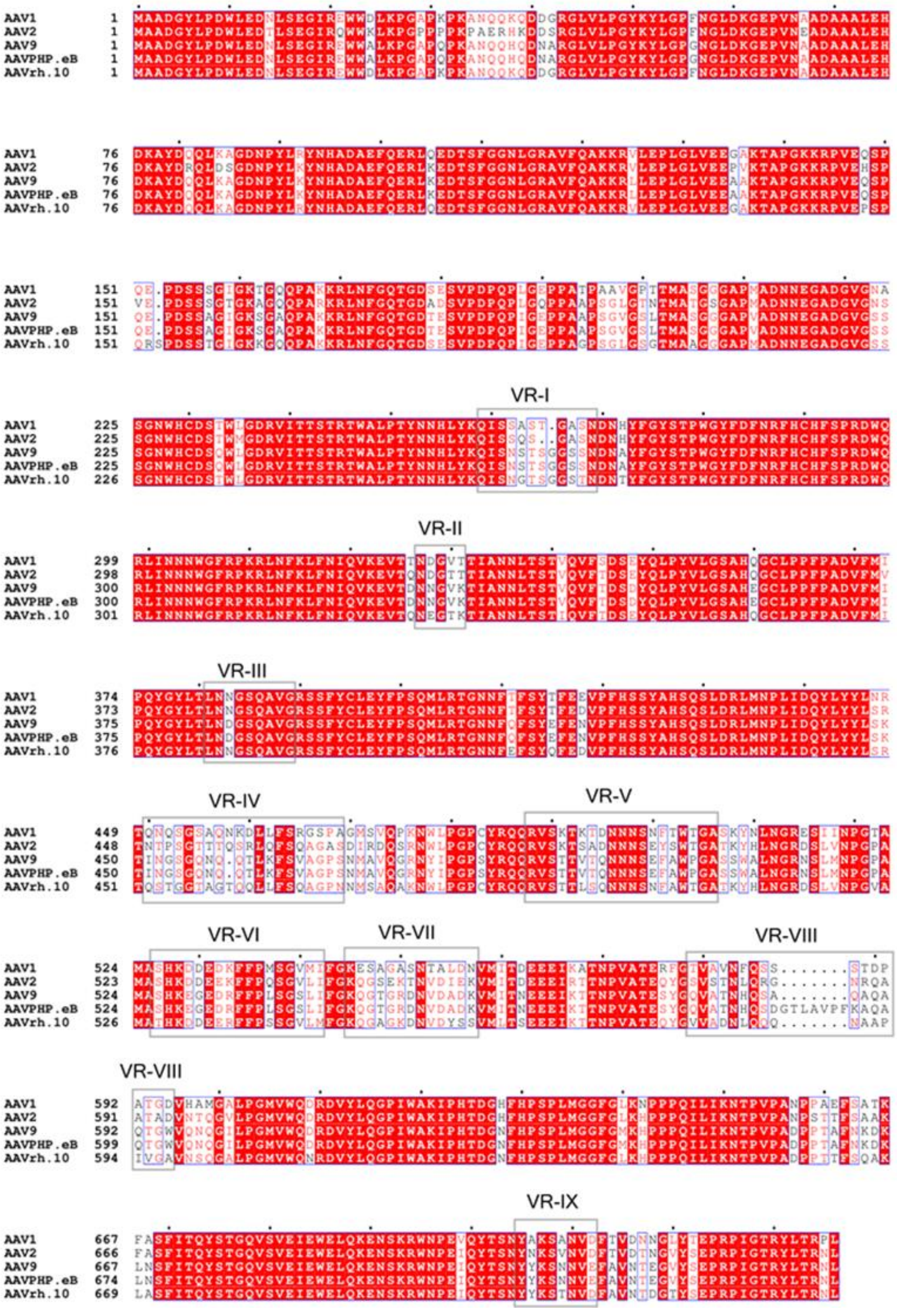

**Supplementary Figure 7.** Amino acid sequence alignment of AAV1, 2, 9, PHP.eB and rh.10 VP1. Sequences were aligned using mafft and displayed with ESPript[1,2]. Grey boxes indicate the designated VRs.

**Supplementary Figure 8**

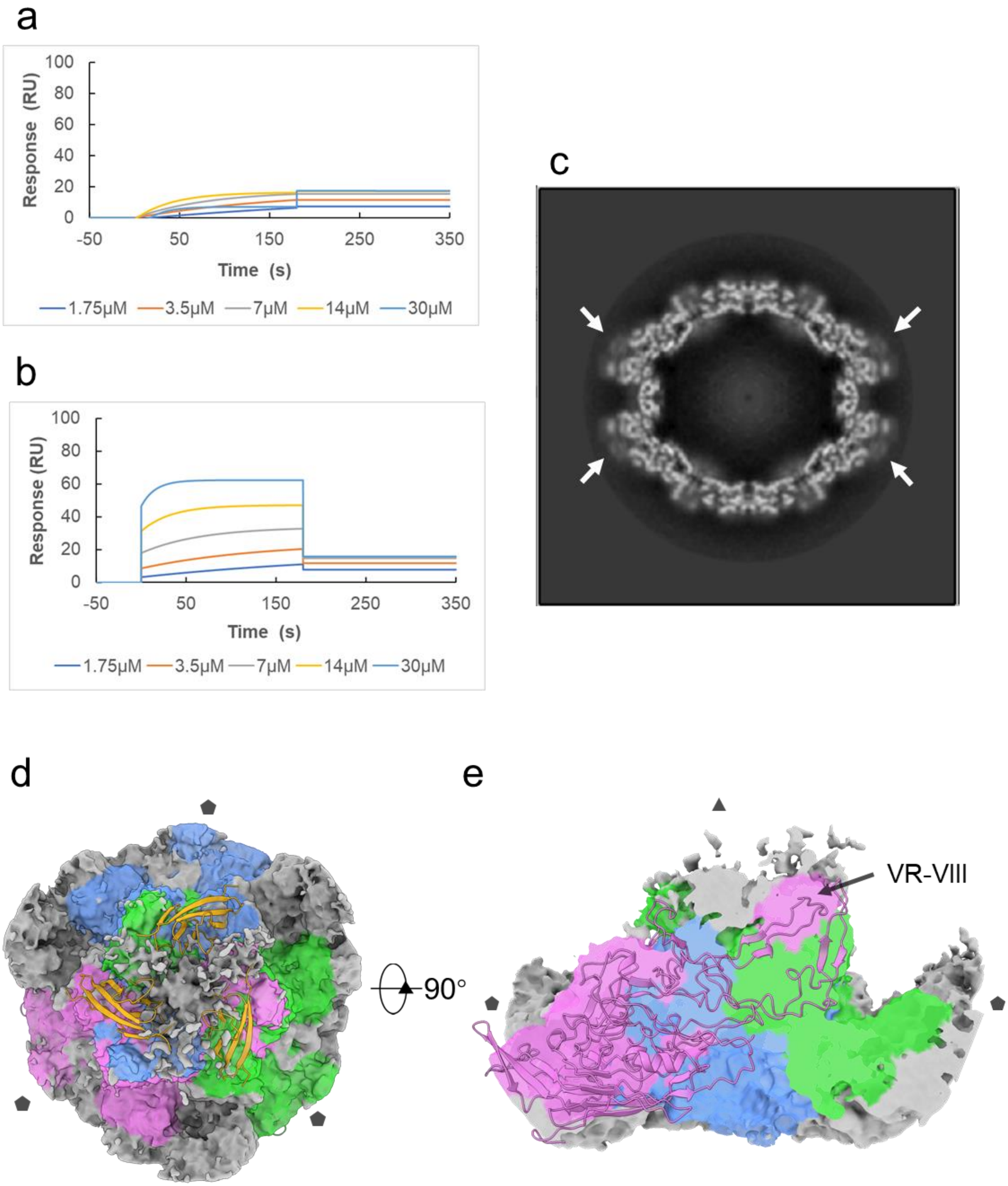

**Supplementary Figure 8.** SPR sensorgrams for binding of LY6A as analyte to immobilized (a) AAV9 and (b) AAVPHP.eB on CM5 chip. (c) central cross-sections through the cryo-EM map of AAVPHP.eB-LY6A. Arrows indicate the additional density above 3-fold axis. (d) Top view of the AAVPHP.eB-LY6A complex 3-fold axis density. A AAVPHP.eB-AAVR trimer model is fitted into AAVPHP.eB capsid density. A 5.5 Å threshold is used to color density around AAVPHP.eB capsomer A, B and C into blue, green and pink, respectively. AAVR PKD2 is presented by ribbon diagrams in gold. (e) is the 90° around X axis rotation and side view section of (d), a capsomer C ribbon diagram (pink) is placed into the corresponding position in density.

**Supplementary Figure 9**

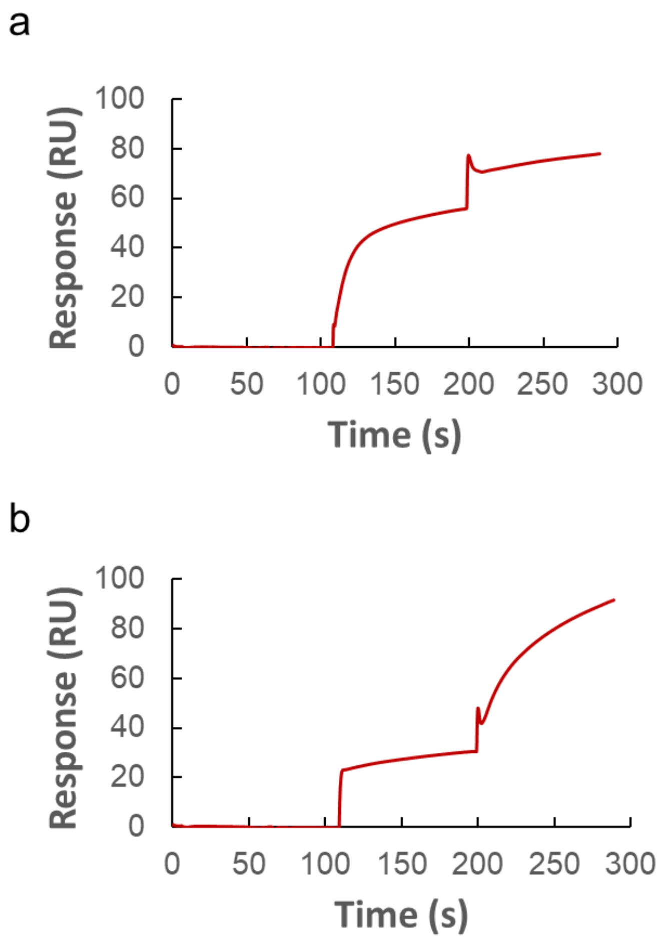

**Supplementary Figure 9.** Binding competition assay performed with SPR. Biacore S200 A-B-A injection method was used with (a) 90 s injections of AAVR to ensure saturation or near-saturation was reached prior to injection of LY6A,and with (b) 90 s injections of LY6A to ensure saturation or near-saturation was reached prior to injection of AAVR.

### Supplementary Tables

#### Supplementary Table 1. Cryo-EM data collection, refinement and validation statistics

| | AAV9 | AAV9 -AAVR | AAVPHP.eB | AAVPHP.eB -AAVR |
|---|---|---|---|---|
| **Data collection and processing** | | | | |
| Magnification | 110,000 | 110,000 | 110,000 | 110,000 |
| Voltage (kV) | 200 | 200 | 200 | 200 |
| Electron exposure (e–/Å$^2$) | 35.7 | 35.7 | 35.7 | 35.7 |
| Defocus range (μm) | -2.0 to -0.7 | -2.0 to -0.7 | -2.0 to -0.7 | -2.0 to -0.7 |
| Pixel size (Å) | 0.93 | 0.93 | 0.93 | 0.93 |
| Symmetry imposed | I1 | I1 | I1 | I1 |
| Initial particle images (no.) | 25,748 | 90,205 | 33,760 | 67,948 |
| Final particle images (no.) | 24,820 | 12,461 | 17,992 | 16,526 |
| Map resolution (Å) | 3.87 | 3.23 | 2.85 | 3.76 |
| FSC threshold | 0.143 | 0.143 | 0.143 | 0.143 |
| Map resolution range (Å) | | | | |
| **Refinement** | | | | |
| Initial model used (PDB) | 3UX1 | 3UX1 | 3UX1 | 3UX1 |
| Model resolution (Å) | | | | |
| FSC threshold | 0.143 | 0.143 | 0.143 | 0.143 |
| Model resolution range (Å) | | | | |
| Map sharpening *B* factor (Å$^2$) | -132.4 | -159.3 | -148.00 | -152.8 |
| Model composition | | | | |
| Non-hydrogen atoms | 4,132 | 4,880 | 4,184 | 4,912 |
| Protein residues | 518 | 615 | 525 | 620 |
| Ligands | 0 | 0 | 0 | 0 |
| *B* factors (Å$^2$) | | | | |
| Protein | 41.68 | 16.33 | 27.76 | 22.75 |
| Ligand | -- | -- | -- | -- |
| R.m.s. deviations | | | | |
| Bond lengths (Å) | 0.006 | 0.007 | 0.009 | 0.011 |
| Bond angles (°) | 0.659 | 0.665 | 0.794 | 0.971 |
| Validation | | | | |
| MolProbity score | 1.91 | 1.80 | 1.94 | 1.79 |
| Clashscore | 7.98 | 7.46 | 5.91 | 5.90 |
| Rotamers outliers (%) | 0.00 | 0.00 | 0.65 | 0.56 |
| Ramachandran plot | | | | |
| Favored (%) | 92.44 | 94.27 | 94.07 | 89.29 |
| Allowed (%) | 7.17 | 5.73 | 5.93 | 10.71 |
| Disallowed (%) | 0.39 | 0.00 | 0.00 | 0.00 |

**Supplementary Table 3. Interaction between AAV9 and PKD2**

| AAVR Residues | Contacts[1] | AAV9 Residues |
|---|---|---|
| R406 | 6 | S263 |
| S413 | 1 | E500 |
| F416 | 2 | S454 |
| I419 | 2 | Q590 |
| T423 | 1 | Q588 |
| S425 | 6 | Q590 |
| T426 | 4 | Q590 |
| D429 | 2 | W503 |
| S431 | 2, 2 | S269, W503 |
| S433 | 2 | S269 |
| T434 | 3, 9, 4 | G267, S268, S269 |
| D435 | 3, 5 | G267, S268 |
| D436 | 3, 3 | N262, S263, S386 |
| D437 | 1, 1, 2, 8, 3 | N262, D384, G385, S386, Q387 |
| K438 | 3, 6, 2 | N270, D384, G385 |
| I439 | 8 | N270 |
| Y442 | 3 | N270 |
| I462 | 2 | W503 |
| K464 | 1 | T590 |

[1]Numbers represent the number of atom-to-atom contacts between the AAVR PKD2 residues and the AAV9 residues, analyzed by the Contact program in the CCP4 suite (with a distance cutoff of 4 Å). Residues colored in blue indicate potential hydrogen bond in-between, residues colored in red indicate potential salt bridge in-between.

**Supplementary Table 3. Interaction between AAVPHP.eB and PKD2**

| AAVR Residues | Contacts[1] | AAVPHP.eB Residues |
|---|---|---|
| R406 | 4 | S263 |
| P414 | 2, 1 | E500, W503 |
| 416F | 9 | S454 |
| Q417 | 1 | S454 |
| E418 | 6 | Q456 |
| I419 | 1 | Q597 |
| T423 | 1, 3 | F594, K595 |
| S425 | 4 | Q597 |
| T426 | 3 | Q597 |
| D429 | 5 | W503 |
| S431 | 3, 1 | S269, D271 |
| Q432 | 3 | S269 |
| S433 | 2, 4 | S268, S269 |
| T434 | 1, 3, 4 | G266, G267, S268 |
| D435 | 3, 9 | N262, G267, S268 |
| D436 | 4, 8, 1, 1 | N262, S263, S386, Q387 |
| D437 | 1, 2, 4 | D384, G385, S386, Q387 |
| K438 | 6 | N270 |
| Y442 | 1 | N270 |
| I462 | 1 | W503 |

[1]Numbers represent the number of atom-to-atom contacts between the AAVR PKD2 residues and the AAVPHP.eB residues, analyzed by the Contact program in the CCP4 suite (with a distance cutoff of 4 Å). Residues colored in blue indicate potential hydrogen bond in-between
